# Single generation allele introgression into pure chicken breeds using Sire Dam Surrogate (SDS) mating

**DOI:** 10.1101/2020.09.03.281204

**Authors:** Maeve Ballantyne, Mark Woodcock, Dadakhalandar Doddamani, Tuanjun Hu, Lorna Taylor, Rachel Hawken, Mike J. McGrew

## Abstract

Poultry is the most abundant livestock species with over 60 billion chickens raised globally per year. While most chicken are produced from highly selected commercial flocks the many indigenous chicken breeds, which have low productivity and have not been highly selected, play an important role in rural economies across the world as they are well adapted to local environmental and scavenging conditions. The ability to rapidly transfer genetic changes between breeds of chicken will permit the transfer of beneficial alleles between poultry breeds as well as allow validation of genetic variants responsible for different phenotypic traits. Here, we generate a novel inducibly sterile surrogate host chicken. Introducing donor genome edited primordial cells into the sterile male and female host embryos produces chicken carrying only exogenous germ cells. Subsequent direct mating of the surrogate hosts, Sire Dam Surrogate (SDS) mating, recreates pure chicken breeds carrying the edited allele in heterozygous or homozygous states. We demonstrate the transfer and validation of two feather trait alleles, Dominant white and Frizzle traits into two pure chicken breeds using the SDS surrogate hosts. This technology will allow the rapid reconstitution of chicken breeds carrying desired genetic changes to investigate climate adaptation and disease resilience traits.

## Main

Poultry is the most abundant livestock species with over 60 billion chickens or 8 chickens raised for each human on the planet each year^1^. Commercial hybrid lines, which have been developed over many decades of selective breeding for meat or egg production, are most common and are extremely productive under controlled diet and environmental conditions. Local populations of chicken that are genotypically, phenotypically, and geographically distinct from each other hold nutritional, economic and cultural importance for smallholder farmers. Although their productivity is much lower than commercial lines, they are posited to be adapted to local climatic and pathogenic insults^2,3^.

Globally, almost 1600 distinct regional breeds of chicken are recognised^4^ and the utilisation of this diversity through the identification and validation of genetic variants for increased resistance to disease and heat stress will benefit both commercial and small holder farmers^5^. The ability to efficiently transfer existing and introduce novel beneficial alleles into both commercial and traditional chicken breeds will aid ongoing efforts to validate disease, production, and climatic adaptation traits in chicken for sustainable food production reviewed in^6^.

Primordial germ cells (PGCs) are the lineage restricted stem cell population for the gametes (sperm and eggs) of the adult animal. The chicken is one of the few species in which the PGCs can be isolated from the developing embryo and then propagated *in vitro*^7,8^. *In vitro* propagated PGCs can be used for the editing of the chicken genome and the cryopreservation of chicken breeds^9–13^. As the chicken develops in a laid egg, PGCs can easily be microinjected into surrogate host chicken embryos, which are subsequently hatched, raised and bred to produce offspring in which one-half of their chromosomes derive from the donor PGCs. A limitation of this strategy is that the gonads of host embryos contain both introduced ‘donor’ PGCs as well as the endogenous ‘host’ PGCs, reducing the chance that offspring from subsequent matings will be derived from ‘donor’ PGCs. To improve the transmission of donor genetic material, it is advantageous to reduce or eliminate the host’s own PGCs. A sterile female surrogate host, a ‘dam’ line containing a knockout of the *cDDX4* gene, was recently shown to lay eggs solely deriving from introduced ‘donor’ female PGCs isolated from an independent breed of chicken^14,15^. The mating sterile male and female (sire and dam) surrogate hosts would permit the direct reconstitution of a pure chicken breed from exogenous PGCs and the generation of homozygous offspring for an introduced genomic variation.

### iCaspase9 expressed from the DAZL locus selectively ablates the germ cell lineage in birds

We aimed to produce a surrogate host chicken line in which the germ cell lineage of both males and females could be conditionally ablated. The inducible caspase 9 protein consists of a truncated human Caspase 9 protein fused to the FK506 binding protein (FKBP) drug dependent dimerization domain^16^. The chemical compound, AP20187 (B/B), induces the dimerisation of FKBP and subsequent activation of the adjoining caspase 9 protein leading to induced apoptotic cell death. iCaspase9 has been previously used as a cellular suicide gene for human stem cell therapy^16–18^ and to ablate cell lineages in transgenic animals^19,20^. We also produced a transgene containing equivalent region of the chicken caspase 9 protein, aviCaspase9 (Supplementary Materials and Methods). To specifically drive the expression of iCaspase9 gene in the germ cell lineage, we used CRISPR/Cas9 mediated homology directed repair (HDR) to target the iCaspase9 construct after the last coding exon of the *cDAZL* gene in *in vitro* propagated chicken PGCs (Fig. 1a). The *cDAZL* gene is highly expressed in migratory PGCs and the embryonic gonad^21,22^. The iCaspase9 transgene was preceded by a 2A peptide sequence and followed by a second 2A peptide and a GFP reporter gene to mark cellular expression. Female PGCs were transfected with the constructs and GFP-expressing PGCs were purified by flow cytometry four weeks post transfection. Targeted GFP^+^ PGCs were assayed for proliferation after addition of the B/B dimerization compound. The growth of PGCs containing either the iCaspase9 or aviCaspase9 transgene was inhibited at nanomolar concentrations of the B/B compound compared with control PGCs targeted with a GFP reporter alone (Fig. 1b).

**Fig. 1:**
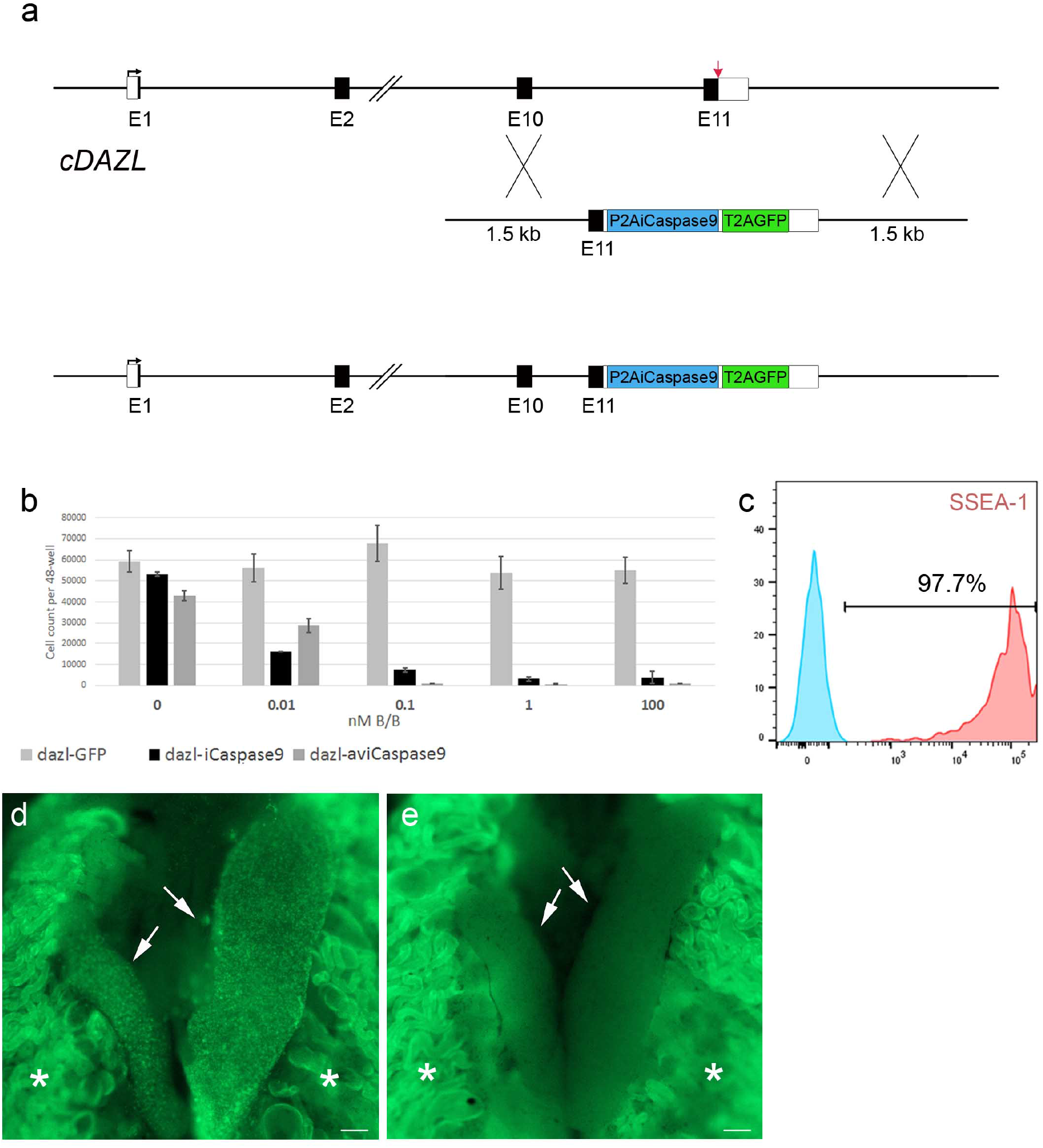
iCaspase9 mediated ablation of the germ cell lineage. **a,** CRISPR/Cas9 mediated recombination of an iCaspase9 and GFP reporter gene into the final coding exon of the *cDAZL* locus. red arrow, guide target **b,** 500 PGCs targeted with a human iCaspase9 or a chicken iCaspase9 (aviCaspase9) transgene were cultured in the presence of different concentrations of the B/B dimerization compound for 10 days. n = 6 for each data point. **c,** 10day gonads were examined for co-expression of GFP and SSEA-1; blue, no primary control. **d,** control and (**e**) B/B treated iCaspase9 G2 embryos imaged day 10 of incubation. Arrows indicate the embryonic gonads. *, autofluorescence in the underlying mesonephros.

GFP^+^ sorted female PGCs were injected into the embryonic vascular system of HH stage 16^+^ (embryonic day 2.5) embryos of the *cDDX4* knockout chicken line. Female chicken ablated for *cDDX4* contain no oocytes post hatch and will produce offspring solely deriving from introduced female PGCs^15^. The injected embryos were sealed and incubated to hatch, then raised to sexual maturity. The *cDDX4* female surrogate hosts were bred to wildtype cockerels and the G_1_ offspring were screened for the presence of the iCaspase9 transgene. 22% of the G_1_ offspring contained the Caspase9 transgene (n= 8 of 74) (Supplementary Table 1 and 2, Supplementary Fig. 1). iCaspase9 and aviCaspase9 G_1_ cockerels were mated to wildtype hens and the G_2_ embryos were examined for GFP expression. GFP fluorescence was observed in the gonads of developing male and female embryos PCR positive for the transgene (Fig. 1d; Supplementary Fig. 3). GFP^+^ cells were positive for the germ cell markers SSEA1 and DDX4, indicating the transgene was expressed in the germ cell lineage (Fig. 1c, Supplementary Fig. 4). Different concentrations of B/B dimerization drug were delivered to chicken embryos *in ovo* which were incubated and examined for both GFP expression and germ cells. We observed that the iCaspase9 gene was more highly expressed in the gonadal germ cells (Supplementary Fig. 4) and more effective for *in ovo* germ cell ablation than the aviCaspase 9 transgene (Fig. 1 d,e, Supplementary Fig. 5-7).

### Removal of the Dominant white mutation from a white feathered chicken breed

The plumage colour of local indigenous chicken populations show great regional diversity and smallholder farmers value coloured plumage for aesthetic, cultural and economic reasons^23,24^. The Dominant white (DOW) feather trait is fixed in the most common commercial layer chicken breed, the White Leghorn (WL), giving them their characteristic pure white plumage^25^. The dominant white (DOW) allele (*I*) has been identified as a putative 9 bp insertion within exon 10 of the *cPMEL17* gene, which creates a 3 amino acid (WAP) insertion in the transmembrane region of the cPMEL17 protein^26^. PMEL17 is a melanocyte transmembrane glycoprotein involved in eumelanin deposition and fibril formation in melanosomes^27^.

The removal of the DOW allele 9 bp insertion from *cPMEL17* in a WL chicken is expected to restore the underlying feather plumage colour in homozygous edited birds. We chose to address this hypothesis using the inbred MHC congenic chicken (line 6) research line of WL chicken. Sequencing of line 6 chickens revealed that this line is homozygous for the DOW allele and Sex-linked barring B2 allele, which generates a barred plumage in female chicken (W/B2) and white feathered males (B2/B2) (Supplementary Figure 8)^28^. Thus, removal of the DOW mutation from line 6 WL chicken should, in principle, generate barred feathered females and white feathered males.

To selectively remove the DOW mutation, we propagated PGCs from single male and female line 6 stage 16^+^ HH (day 2.5) embryos^8^. A CRISPR guide along with a high fidelity Cas9 (SpCas9-HF1) and an ssODN donor template containing the wildtype *cPMEL17* sequence, were transfected into the PGC cultures. Distinct 95 bp ssODN donor templates were used for transfection of the female and male PGC cultures with the purpose of tracking the parental derived alleles in the G_1_ generation. These templates differed by a single synonymous nucleotide change which was introduced into the female PGCs, whereas the wildtype *cPMEL17* nucleotide sequence was introduced into the male PGCs (see Supplementary Materials and Methods for ssODN sequences). After clonal isolation and culture, approximately 50% of PGCs clones were identified to contain an edited wildtype allele (*cPMEL17^-WAP^*) in both male and female PGC lines. Unexpectedly, the second allele in most male and female clones containing an identical 15 nucleotide deletion adjacent to the Cas9 cut site (Fig. 2a). This in-frame deletion removed the guide binding site and created a 5 amino acid deletion in the transmembrane domain whilst leaving the PAM sequence and the DOW insertion allele intact (*cPMEL17^Del+WAP^*). This deletion was predicted to occur at this Cas9 cleavage site by micro-homology mediated repair predictors^29,30^.

**Fig. 2:**
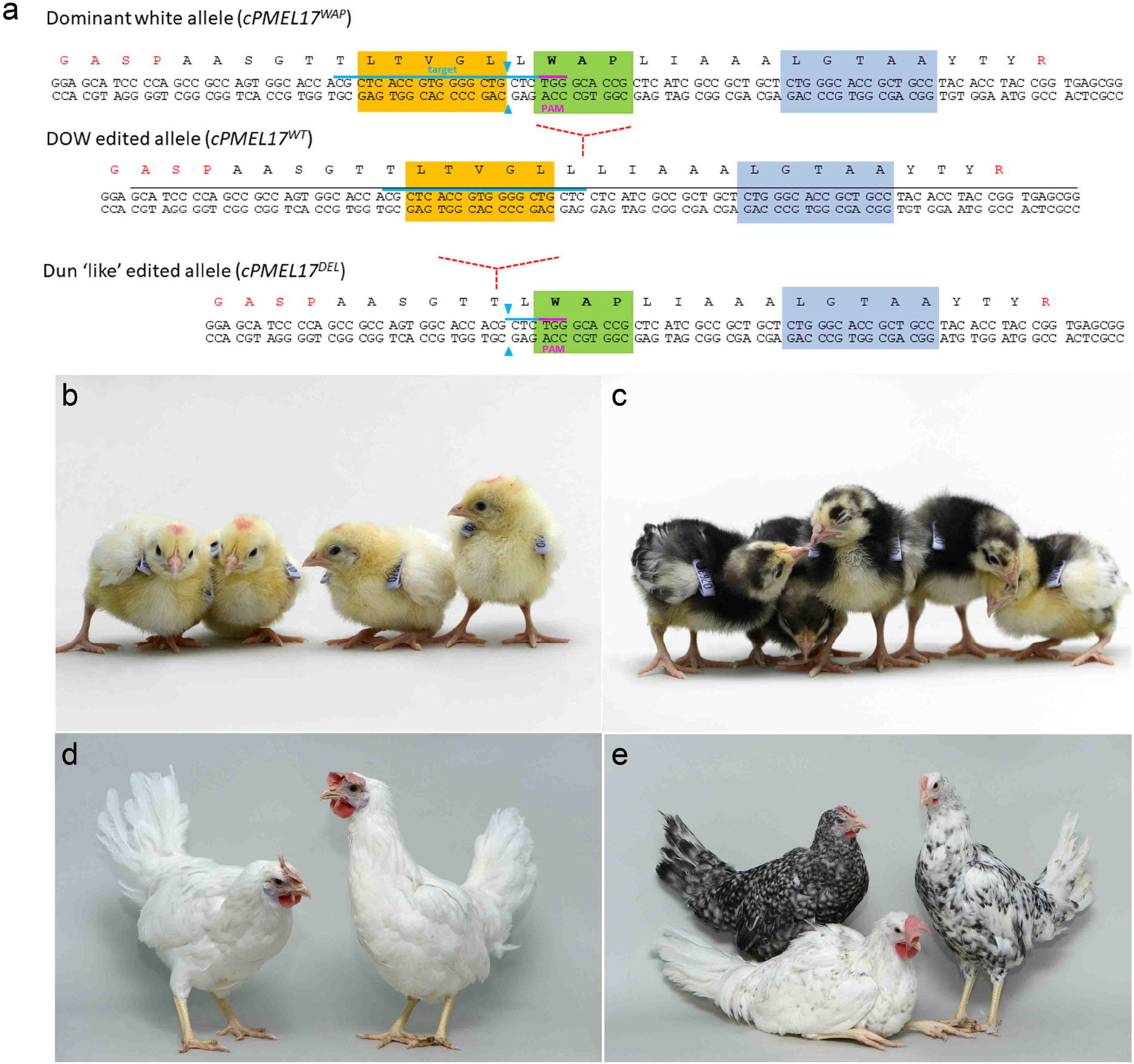
Genome edit of the Dominant white locus from a white leghorn chicken breed. **a,** Locus containing WAP insertion in the 10^th^ exon and the transmembrane domain of *cPMEL17*. Genome edited *PMEL17* locus to remove WAP allele and creating a 5 amino acid deletion allele. **b-d,** wildtype Line 6 chicks and adults. **c-e,** *PMEL17-edited* chicks and adults; females (standing) were barred and speckled feathers, males (sitting) were white with black dots.

We used these PGCs to generate a *cPMEL17* allelic series containing the two edited changes. To directly produce genome edited G_1_ chicks, we mixed edited female and male PGCs and coinjected these with B/B dimerization compound into the embryonic vascular system of HH stage 16^+^ iCaspase9 and aviCaspase9 chicken embryos in windowed eggs (Fig. 5). The host shells were sealed and incubated until hatching (Supplementary Table 1). The Caspase9 surrogate host chicks were raised to sexual maturity and directly mated to each other in natural mating groups; a process we call Sire Dam Surrogate (SDS) mating. Three independent mating groups were produced and fertile eggs were incubated and hatched (Table 1). All hatchlings from the iCaspase9 mating groups were seen to display a black plumage pigmentation. In addition, a PCR analysis did not detect the iCaspase9 transgene in any of the offspring (Fig. 2c, Table 1). When mature, the cockerels displayed a floppy comb that is consistent for Line 6 birds (Fig. 2). Furthermore, a principle component (PC) analysis indicated that all offspring from the iCaspase9 clustered with control Line 6 birds (Fig. 3). In contrast, the aviCaspase9 surrogate host group produced many chicks with yellow feathers and a PCR analysis of the offspring detected the iCaspase9 transgene indicating the endogenous germ cells were not completely ablated in this host (Table 1). The egg lay rate and fertility from natural matings for the iCaspase9 mating groups was high and similar to that observed for the aviCaspase9 mating group. However, the hatchability from the iCaspase9 surrogate host mating groups was lower than expected (~60%) in comparison to the aviCaspase9 group (82%). This suggests that the *in vitro* culture/manipulations of the PGCs may have lowered overall hatchability. However, survivability of the Line 6 hatched chicks was not different from control Line 6 birds (data not shown).

**Fig. 3:**
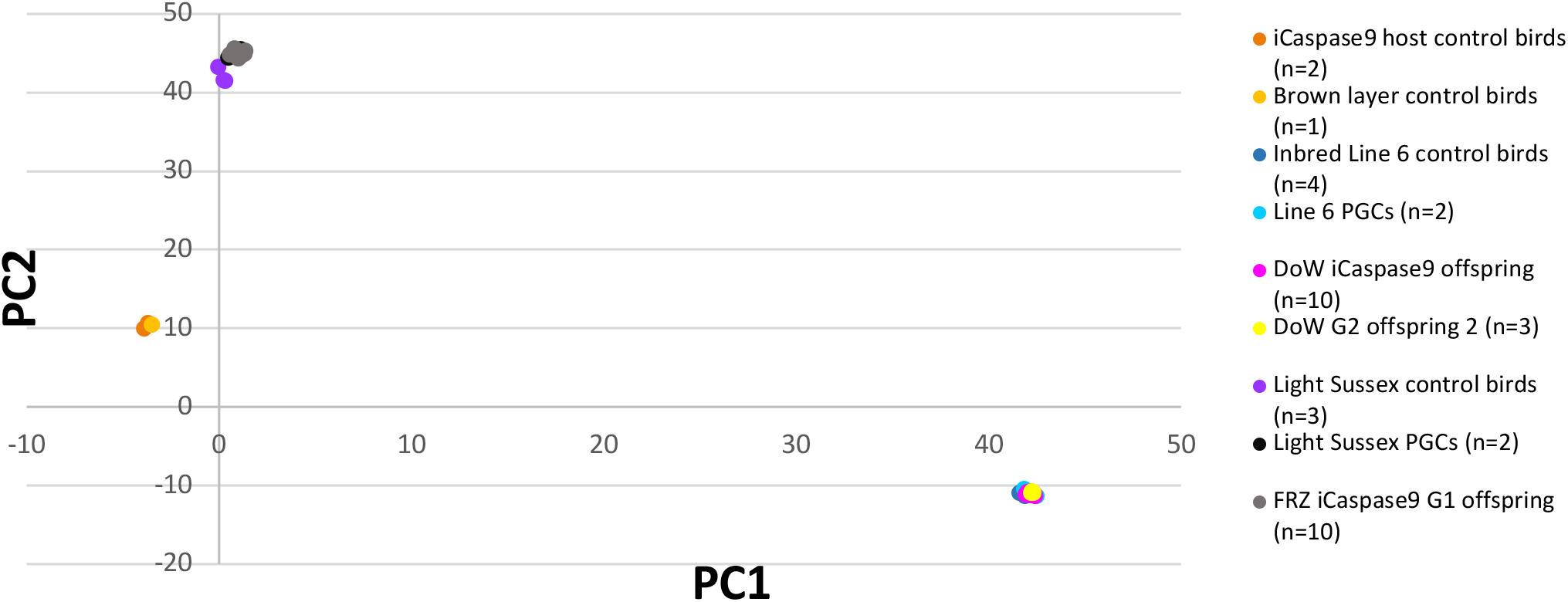
Principle component analysis of G_1_ offspring from iCaspase9 surrogate host birds. DNA samples were genotyped on a 66k SNP chip and analysed for PCs. Three chicken breeds were analysed: the Hy-Line brown layer line and iCaspase9 surrogate hosts, the Light Sussex breed, and Line 6 WL controls. Offspring from iCaspase9 hosts injected with LSX PGCs (grey) clustered with Light Sussex controls. Offspring from iCaspase9 surrogate hosts injected with DOW PGCs (pink) clustered with Line 6 controls.

**Fig. 4:**
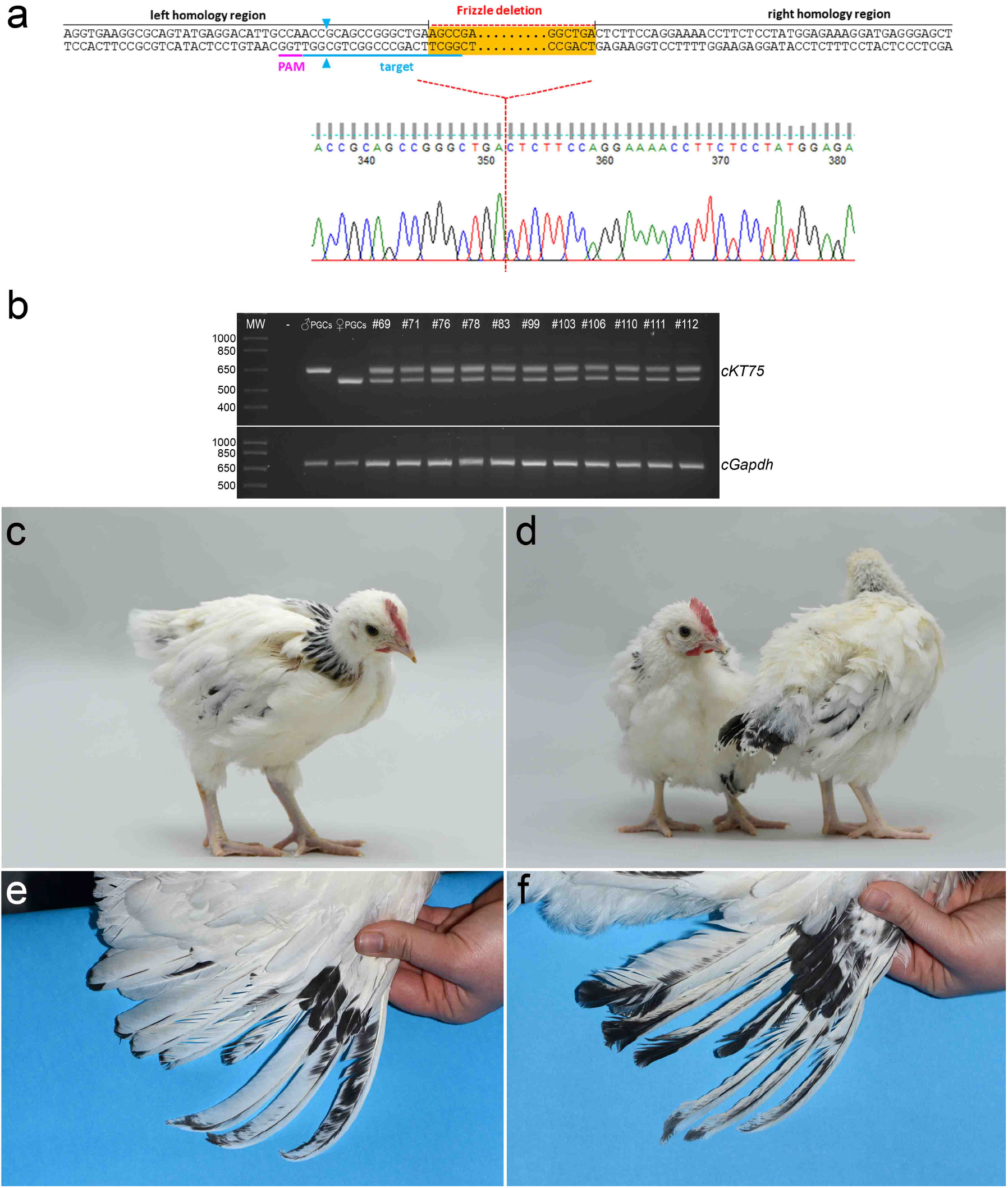
Genome edit to introduce the Frizzle mutation into the Light Sussex chicken breed. **a,** A single guide (blue) targets cleavage 12bp from the intended deletion was transfected with a ssODN containing 50bp homology on each side of the deletion. Sanger sequencing traces shows the biallelic deletion in female LSX PGCs. **b,** PCR analysis of LSX edited PGCs and G_1_ offspring from iCaspase9 hosts. The wildtype locus produces a PCR product of 657 bp. The edited locus produced a PCR product of 573 bp. **c,e,** Control 6 week old offspring and wing (18 weeks) from LSX birds. n =10/10 female birds. **d,f,** FRZ heterozygote LSX G_1_ offspring (6 weeks) and wing (18 weeks) displaying crumpled flight feathers, n = 5 of 5 female birds.

**Fig. 5:**
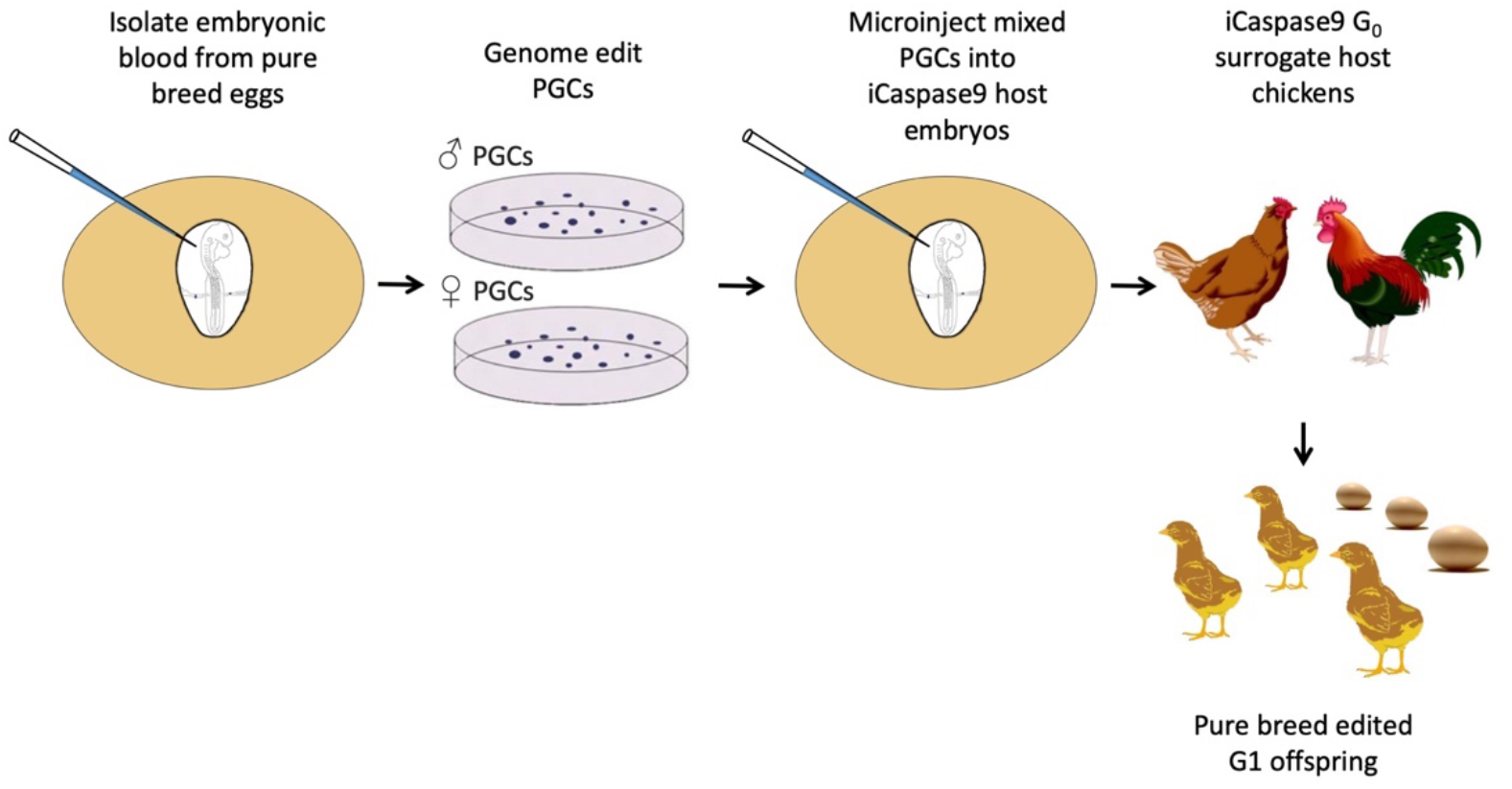
Schematic of SDS mating. PGCs are isolated and propagated *in vitro* from a pure chicken breed. After genome editing and clonal isolation, the male and female PGCs are mixed and injected into iCaspase9 surrogate host embryos. The embryos are hatched, raised to sexual maturity then mated. Laid eggs and hatched offspring are from the donor breed of interest and contain the desired genome edit.

**Table 1.**
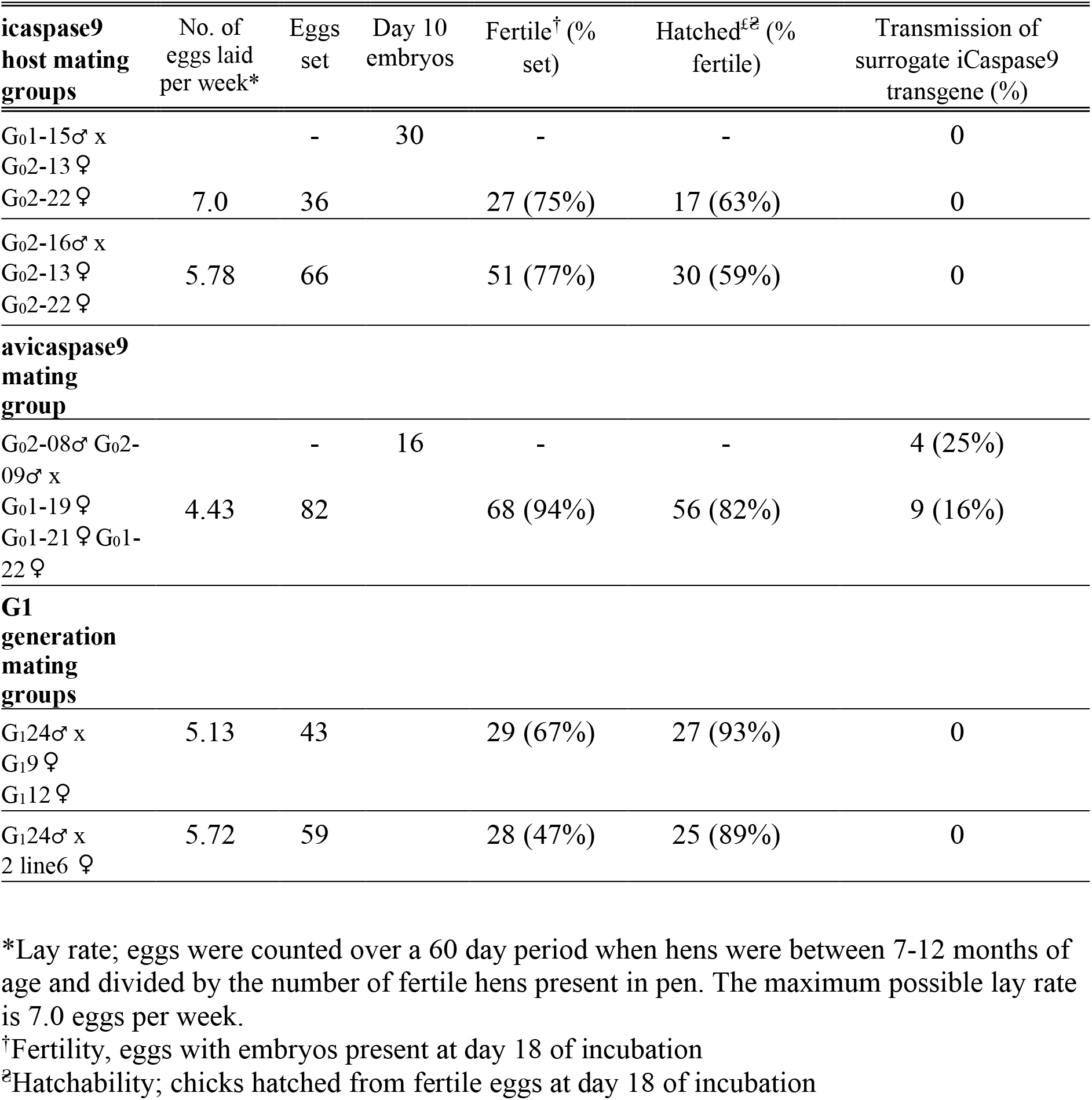
Germline transmission rates from surrogate hosts injected with donor pMel17 edited PGCs

The adult G_1_ birds displayed three colour phenotypes; (Fig. 2e) cockerels were spotted white, and hens had barred or speckled feathers. Sanger sequencing analysis and restriction digest genotyping revealed that the barred/speckled females had the genotypes *cPMEL17^-WAP/-WAP^, cPMEL17^-WAP/Del+WAP^, cPMEL17^Del+WAP/Del+WAP^* (Supplementary Fig. 9 a-c), The *cPMEL17^-WAP/WAP^* birds all contained a single tracking nucleotide indicating that these offspring are descended from one male and one female PGC (Supplementary Fig. 9a). Crossing a *cPMEL17^-WAP/WAP^* G_1_ male and females confirmed three feather colour morphs (cockerels, spotted white; hens, barred or speckled feathers) were present in the G_2_ generation which corresponded with the expected feather colour phenotypes. Breeding *cPMEL17^-WAP/-WAP^* G_1_ males with Line 6 wildtype birds generated all white feather offspring showing that the edited alleles were recessive to the DOW allele (Supplementary Fig. 10). This result suggests the length of the transmembrane domain is important for optimal function. The wildtype cPMEL17 transmembrane domain is 25 amino acids (aa) in length, the WAP insertion increases this length to 28 aa and creates a dominant negative protein. The naturally occurring *Dun* loss of function allele reduces the transmembrane length to 20 aa. Here, we restore the wild-type 25 aa length in the −WAP allele (*cPMEL17^-WAP^*) and create a novel mutation resulting in a 27 aa transmembrane domain, *cPMEL17^Del+WAP^*. Both of which restore eumelanin production as attested to by the dark barred feather female offspring.

### Introduction of the Frizzle (FRZ) feather trait into a traditional European breed

Frizzled feather chicken are highly valued in Western Africa and this trait is posited to confer adaptability to tropical climates^31,32^. The frizzle (FRZ) feather allele (*F*) is a naturally occurring splice variant caused by an 84 bp deletion in the a-keratin 75 gene (*cKRT75*) leading to a curved feather rachis and barbs^33^. Homozygote FRZ birds have severely frizzled and brittle feathers that are easily broken, however heterozygote FRZ chicken display an attractive ruffled, frizzled feather phenotype. An autosomal recessive modifier gene (mf) that lessens the effects of the FRZ mutation is present in many European chicken breeds, leading to a minor frizzled phenotype in young chicken and a crumpling of the barbs of the anterior flight feathers^34,35^. To produce heterozygote frizzle chicken, we first isolated and propagated male and female PGCs from eggs deriving from a flock of Light Sussex chicken. The Light Sussex (LSX) is a traditional dual purpose British breed with black tail feathers and black stripped hackles^36^ (Fig. 4c). We again used high fidelity Cas9, SpCas9-HF1, and a 100 bp ssODN template to introduce the FRZ deletion into female LSX PGCs. After clonal isolation and propagation, we found that 10% of the PGCs clones were homozygous for the FRZ genome edit (Fig. 4a).

To directly produce heterozygote FRZ edited LSX G_1_ birds we used the iCaspase9 surrogate host eggs and SDS mating. Homozygous *cKRT75* edited female LSX PGCs and wildtype male LSX PGCs were mixed and co-injected with the B/B compound into the embryonic vascular system of stage 16^+^ iCaspase9 chicken embryos in windowed eggs. The eggs were sealed and incubated to hatch (Supplementary Table 1). The iCaspase9 surrogate host brown layer G0 birds were raised to sexual maturity and naturally mated. Fertile eggs from the SDS matings of two independent groups were incubated and hatched. Egg laying, fertility and hatchability were appropriate for layer chicken and are shown Table 2. PCR analysis of both embryos and hatched chicks found that all offspring were heterozygote for the frizzle allele indicating that they were derived from a single wildtype male PGC and a single edited female PGC (Fig. 4b). A PC analysis showed that the iCaspase9 offspring clustered with control LSX birds indicating all offspring derived from donor LSX PGCs (Fig. 3). The hatched offspring displayed typical LSX colouration and a feather phenotype representative of a modifier FRZ heterozygote; obvious frizzled feather phenotype at 4 weeks of age and the flight feather were crumpled as adults (Fig. 4c-f).

**Table 2.**
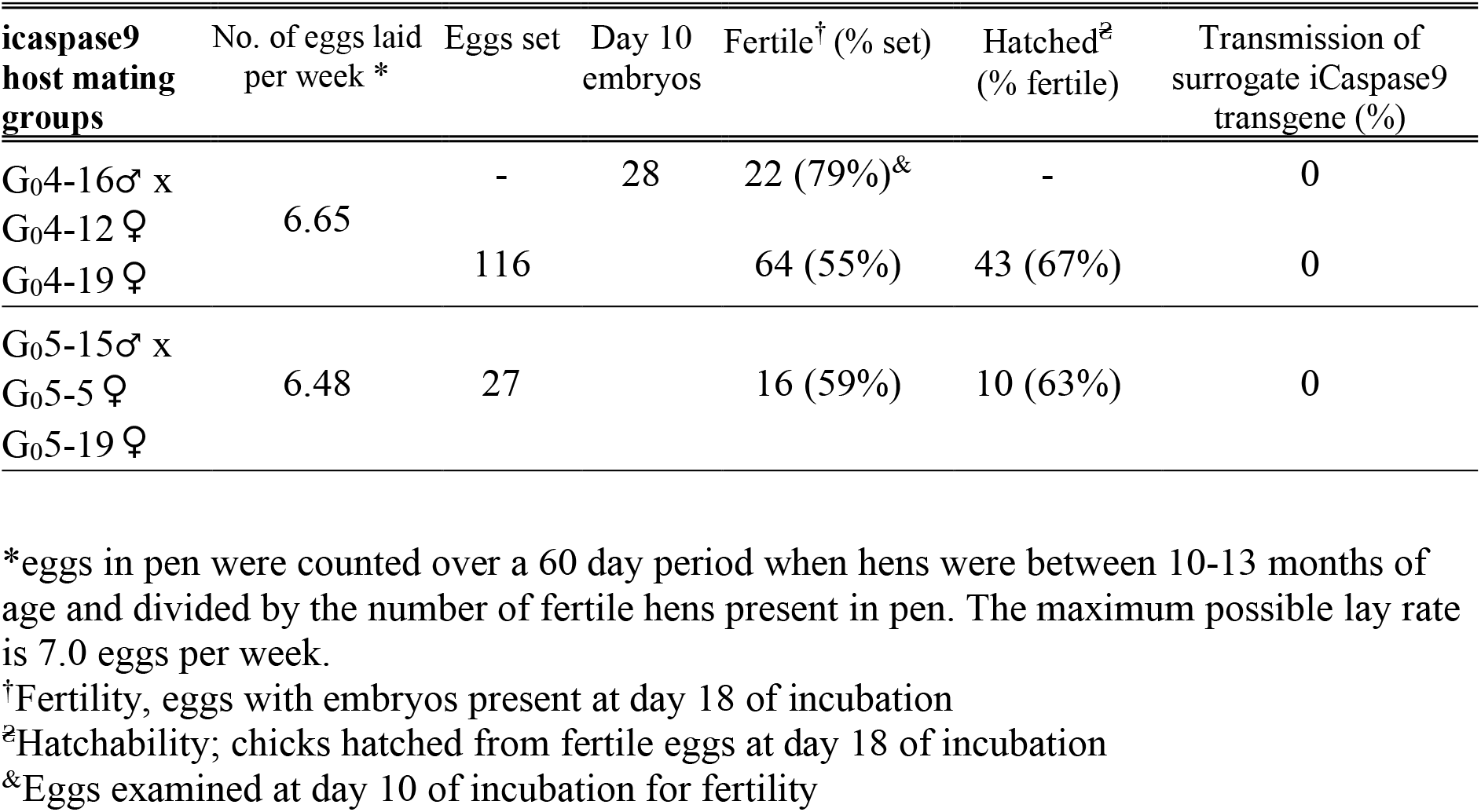
Germline transmission rates from surrogate hosts injected with donor *cKT75* edited ♀ LSX PGCs and wildtype ♂ LSX PGCs

## Discussion

Our results demonstrate the usefulness of sterile surrogate hosts in a bird species for producing pure breed offspring by the direct mating of surrogate sires and dams. As the chicken is a model organism for the study of embryogenesis, neural development, and immunity in bird species, this technique offers a reliable platform for investigating gene function and disease resistance in poultry. As two generations are required to produce the homozygous offspring from genome edited PGCs and the generation time for chicken is 5-6 months, SDS breeding greatly accelerates the production of homozygote offspring for validation of genetic variants. SDS mating is particularly amenable to species in which the embryo develops *ex utero* such as birds, fish, amphibians, and reptilian species^37–40^.

The cryo-conservation of avian species is challenging due to both the yolk filled egg and difficulty in cryopreserving avian sperm. SDS mating allows the regeneration of chicken breeds directly from frozen reproductive material. Indeed, we demonstrated the reconstitution of two pure chicken breeds from cryopreserved PGCs: the White Leghorn and the Light Sussex. Here, we also show that breed regeneration can be coupled with the introduction and removal of genetic traits. Allelic validation may be increased as SDS matings produced large numbers of full siblings deriving from the same male and female PGCs. This could be useful for determining gene function whilst reducing phenotypic variation between offspring. However, chicken flocks are maintained as highly genetically diverse populations to avoid inbreeding which causes dramatic reductions in fertility and hatchability. It will be necessary to multiplex multiple genotypes through single surrogate hosts in order to regenerate outbred populations.

## Materials and methods

### Chicken breeds and welfare

The MHC Line 6 congenic line of white leghorn birds^41^ and the Light Sussex dual purpose chicken breed^36^ are maintained as a closed breeding population of 150 birds in the NARF SPF facility (UK) and provided fertile eggs used for PGC derivations. Individual Line 6 birds were sequenced to confirm the presence of the dominant white allele and the absence of the recessive white allele^42^. The barred feather allele was identified in line 6 by analysis of the *CDKN2a* locus for SNP2 and SNP4^28^. The Caspase9 lines of chickens were generated using a Hy-line Brown layer PGCs. Heterozygous and homozygous cockerels carrying the iCaspase9 transgene were crossed to Hy-line hens to produce fertile eggs for injection and hatching. DDX4 ZZ-heterozygote males were crossed to Hy-line hens to produce fertile eggs for injection. All animal management, maintenance and embryo manipulations were carried out under UK Home Office license and regulations. Experimental protocols and studies were approved by the Roslin Institute Animal Welfare and Ethical Review Board Committee.

### Chicken PGC culture and transfection

PGC lines were derived from the blood of stage 15-16^+^ embryos and propagated in culture as previously described in Whyte *et al*. The embryos and PGC lines were sexed using a W-chromosome specific PCR as previously described in Clinton *et al*. Both male and female PGC cultures were derived from Hy-line, Light Sussex, and Inbred Line 6 (White Leghorn) embryos. Each PGC line was in culture for approximately 3 weeks before freezing in aliquots of 50,000 cells resuspended in 125 μl of Stem-Cellbanker (Amsbio) and stored at −150 °C.

For generation of iCaspase9 birds; 1.0 ug of iCaspase9 or aviCaspase9 targeting vector and 1 ug of Dazl CRISPR/Cas9 vector were transfected into 1.5 x 10^5^ Hy-line PGCs using Lipofectamine 2000 transfection reagent (Thermo Fisher Scientific). PGCs were cultured for three weeks and GFP^+^ PGCs were purified by flow cytometry using a FACS-ARIA gated for GFP florescence. For CRISPR editing of *cKRT75* and *cPMEL17;* 2 x 10^5^ of LSX or Line 6 PGCs were transfected with 1.5 μg of CRISPR/Cas9 vector and 0.5 μg of ssODN donor template. After 24 hours in culture the cells were treated with 0.6 μg/ml puromycin for 48 hours to select for cells transfected with CRISPR vector. After selection PGCs were sorted using the FACS Aria III cell sorter into a 96-well plate at 1 cell per well After culturing for 3 weeks, clonal PGC populations were cryopreserved and used for genomic DNA isolation.

### Production of surrogate host chicks

Targeted male and female PGC lines were thawed from storage at −150 °C and cultured for 5-10 days before injection into stage 15-16^+^ HH surrogate host embryos in windowed eggs. 1.0 μl of PGCs were directly injected into the dorsal aorta and all shells were resealed with medical Leukosilk tape (BSN Medical) before incubating until hatch. For experiments using iCaspase9 surrogate host embryos, 1.0 ul of 25mM B/B (in DMSO) (Takara Bio) was added to 50ul PGC suspension before injection and subsequently 50 μl 300ul P/S (containing 30ul of 0.5mM B/B drug) was pipetted on top of the embryo. Founder males and females were naturally mated in pens to produce G_1_ offspring. All offspring were screened by PCR for the presence of iCaspase9 transgene. See Supplementary Table 1 for details on individual experiments.

## Acknowledgements

We thank the members of the Roslin chicken facility (A. Sherman, M. Hutchison, F. Brain, K. Hogan and F. Thomson) for care and breeding of the chickens, Jun Chen and Brenda Flack (Cobb-Vantress) for the SNP chip analysis, and Megan Davey for critiquing the manuscript. We thank Donald Nkrumah for the guidance to edit the frizzle feather allele and Appolinaire Djikling, Bruce Whitelaw, Steve Kemp and the members of the Centre Management Group of the CTLGH for constructive comments on the project.

This research was funded in part by the Bill & Melinda Gates Foundation and with UK aid from the UK Government’s Department for International Development (Grant Agreement OPP1127286) under the auspices of the Centre for Tropical Livestock Genetics and Health (CTLGH), established jointly by the University of Edinburgh, SRUC (Scotland’s Rural College), and the International Livestock Research Institute. The findings and conclusions contained within are those of the authors and do not necessarily reflect positions or policies of the Bill & Melinda Gates Foundation nor the UK Government. This work was supported by the Institute Strategic Grant Funding from the BBSRC (BB/P0.13732/1 and BB/P013759/1) and Innovate UK Agri-Tech funding (BB/M011895/1).

## Contributions

MB, MW, RH, MJM conceived the project. MB, MW, DD, TH, LT, MM, RH, performed the experiments. All authors analysed the results. MB, MJM, RH wrote the manuscript.

