## Supplementary Tables, Figures, and Methods for "Single generation allele introgression into pure chicken breeds using Sire Dam Surrogate (SDS) mating"

**Supplementary Material**

**Supplementary Table 1.** Injection of surrogate host chicks carrying donor PGCs

| Donor PGC genotypes | PGCs injected per embryo | No. of expts | Sire genotype | Total no. of eggs injected | Embryos at day 18 incubation | Chicks hatched (% hatchability) | Correct genotype host* |
|---|---|---|---|---|---|---|---|
| iCaspase9♀<br>aviCaspase9♀ | 3000 | 1 | $ZZ^{DDX4}$ | 23 | | 12 (%) | 2 (17%) |
| pMel17 Line6♀<br>pMel17 Line6♂ | 6000 | 2 | $Dazl^{iCaspase9}$ | 27 | 18 | | 9 (50%) |
| pMel17 Line6♀<br>pMel17 Line6♂ | 6000 | 2 | $Dazl^{aviCaspase9}$ | 26 | 18 | | 5 |
| cKRT LSX♀<br>LSX♂ | 4000 | 1 | $Dazl^{iCaspase9}$ | 23 | | 7 | 4 |
| cKRT LSX♀<br>LSX♂ | 4000 | 1 | $Dazl^{iCaspase9/iCaspase9}$ | 21? | | 8 | 8 (100%) |

*For *DDX4* hosts, 25% of offspring are $WZ^{DDX4}$. The $Dazl^{iCaspase9}$ line was originally heterozygous and transmitted the transgene to 50% of the offspring when bred to wildtype females. A $Dazl^{iCaspase9/iCaspase9}$ male was produced which was bred to wildtype hens and transmitted the transgene to 100% of the offspring.

**Supplementary Table 2.** Germline transmission rates of *DDX4* surrogate hens carrying donor iCaspase targeted PGCs

| Host genotype | No. of surrogates | No. of eggs laid per week* | No. of eggs incubated | Fertility$ (% eggs incubated) | Hatched† (% hatchability) | No. of transgenic chicks (% Transmission₴) |
|---|---|---|---|---|---|---|
| $Z^{DDX4-}W$ | 2 | 3.33 | 85 | 80 (94%) | 74 (93%) | 8 (22%) |
| ZW | 2 | 6.77 | 135 | 115 (85%) | 93 (81%) | 0 (0%) |

*eggs in pen were counted over a 60 day period when hens were between 7-9 months of age and divided by the number of fertile hens present in pen. The maximum possible lay rate is 7.0 eggs per week.
$embryos present at day 18 incubation of eggs incubated.
†chicks hatched from embryos present at day 18 of incubation.
₴the number icaspase9 chicks per number of hatched chicks equals one half the transmission rate due to meiotic reduction.

**Supplementary Figures**

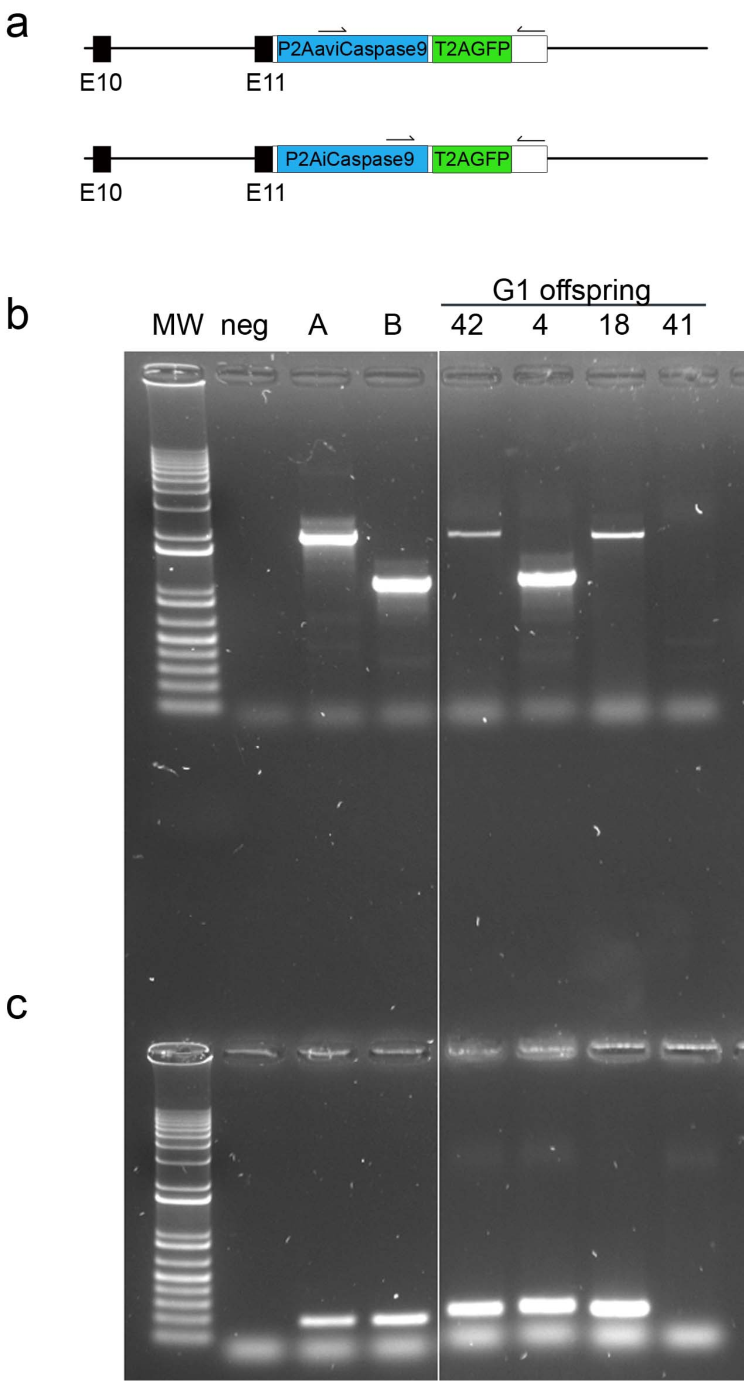

**Supplementary Fig. 1 Generation of iCaspase9 and aviCaspase9 $G_1$ offspring**

**a,** Diagram of targeted *cDazl* locus and primer sites.

**b,** PCR of $G_1$ offspring with Caspase9-specific primers; MW, molecular weight markers, A, targeted PGCs containing aviCaspase9 transgene, B, targeted PGCs containing iCaspase9 transgene. Positive offspring shown here are numbers 42, 18: aviCaspase9; 4: icaspase9. **c,** PCR with primers specific for GFP.

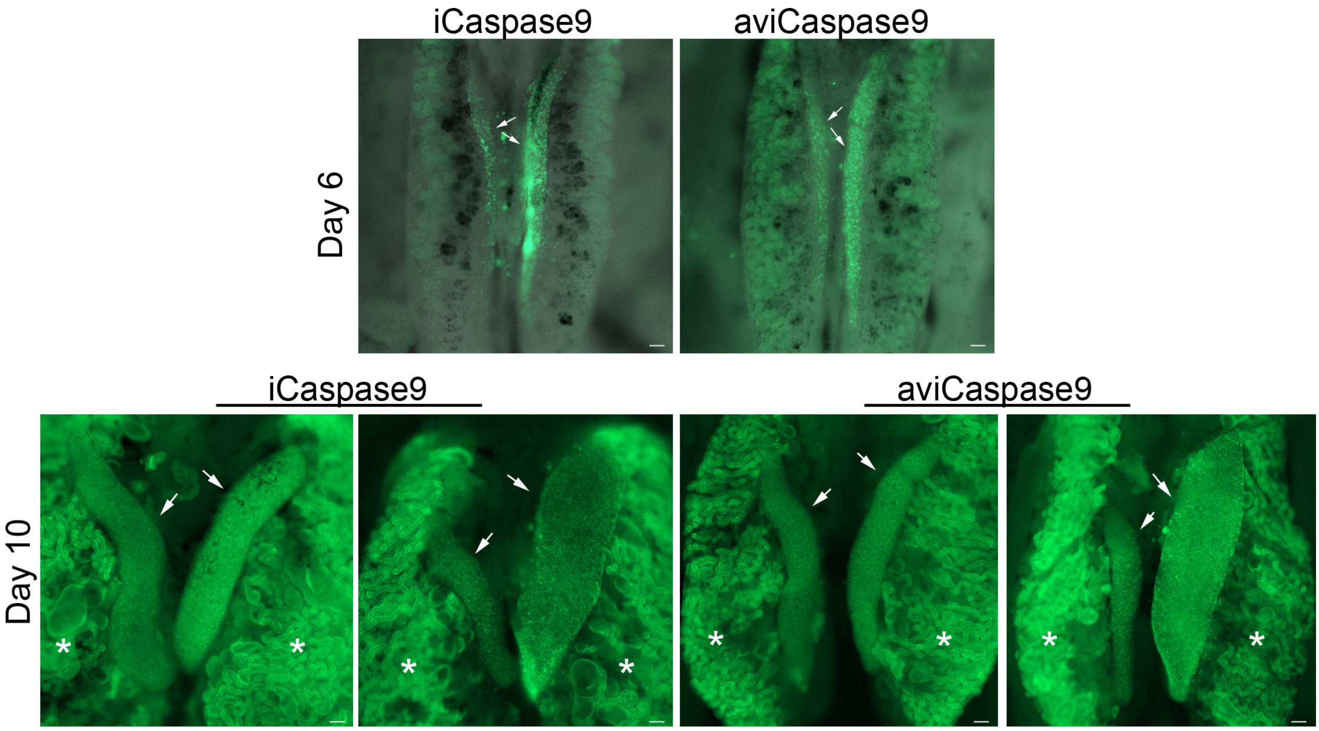

**Supplementary Fig. 2 GFP expression in iCaspas9 and aviCaspase9 G2 embryos $G_2$ chicken embryos**

Representative gonads from day 6 or day 10 incubated $G_2$ embryos that PCR positive for the iCaspase9 and aviCaspase9 genes were isolated and imaged for GFP fluorescence. Arrows indicate the embryonic gonads. *, autofluorescence in the underlying mesonephros. Scale bar, 100 μm.

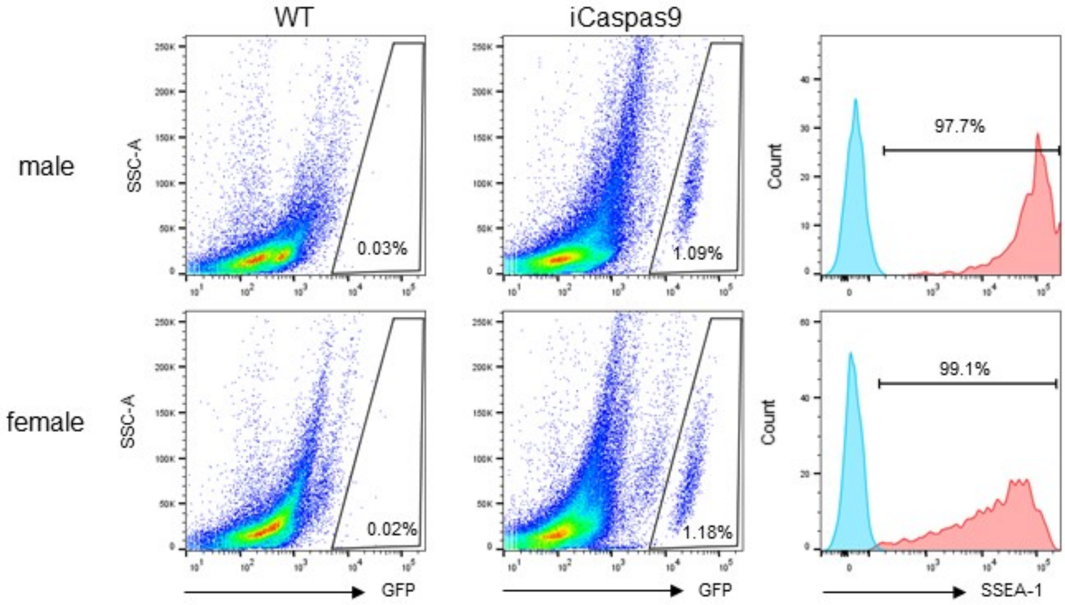

**Supplementary Fig 3. Flow cytometric analysis of gonadal PGCs.**
Gonads from male or female embryos at 10 days of incubation were dissociated and analysed for GFP expression; $GFP^+$ cells were examined for expression of SSEA-1. Co-staining with SSEA-1 antibody (filled red) indicated the entire population of $GFP^+$ cells also express SSEA-1 antigen, as shown in histogram. Blue peak, no primary control.

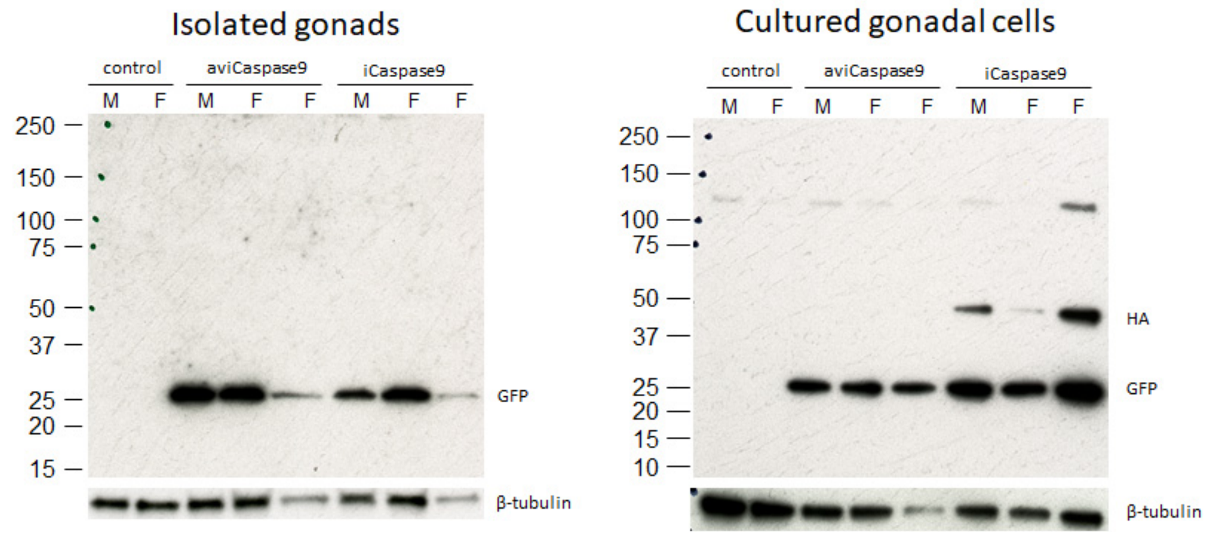

**Supplementary Fig 4. Western Blot analysis of day 14 embryonic gonads**

GFP$^+$ gonads from a single iCaspase9 or aviCaspase9 embryos at 10 days of incubation were examined for GFP or HA-tag expression. Single gonads dissociated and ½ the sample was run per lane (~75mg/lane) or cultured for five days before Western analysis. Control = GFP$^-$ gonads.

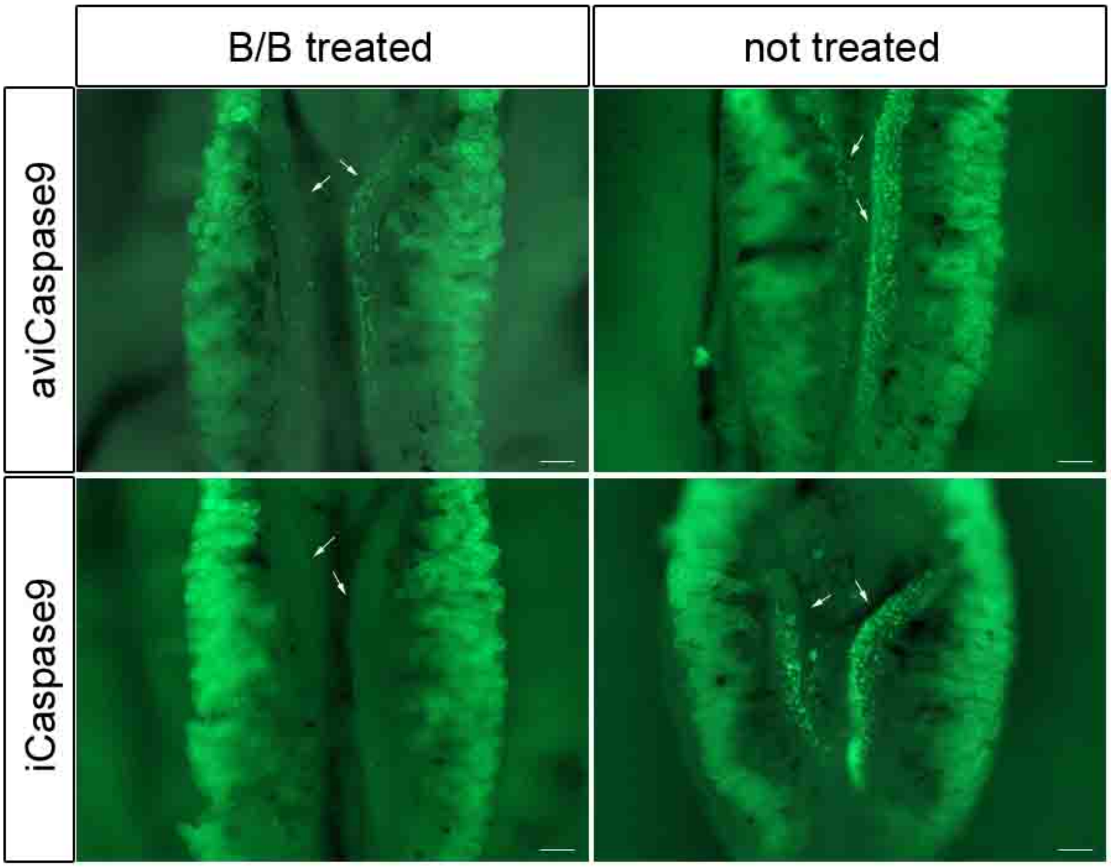

**Supplementary Fig. 5 iCaspase9 embryos are more responsive to B/B than aviCaspase9 embryos.**

Stage 16 HH (day 2.5) embryos were injected with 1ul of 0.1mM B/B into the dorsal aorta and incubated to day 6 of embryonic development. Arrows indicate the $GFP^+$ cells in the gonads. Fewer cells are present in the icaspase9 embryo (n= 3, aviCaspase9; n = 5, iCaspase9). Scale bar, 100 µm.

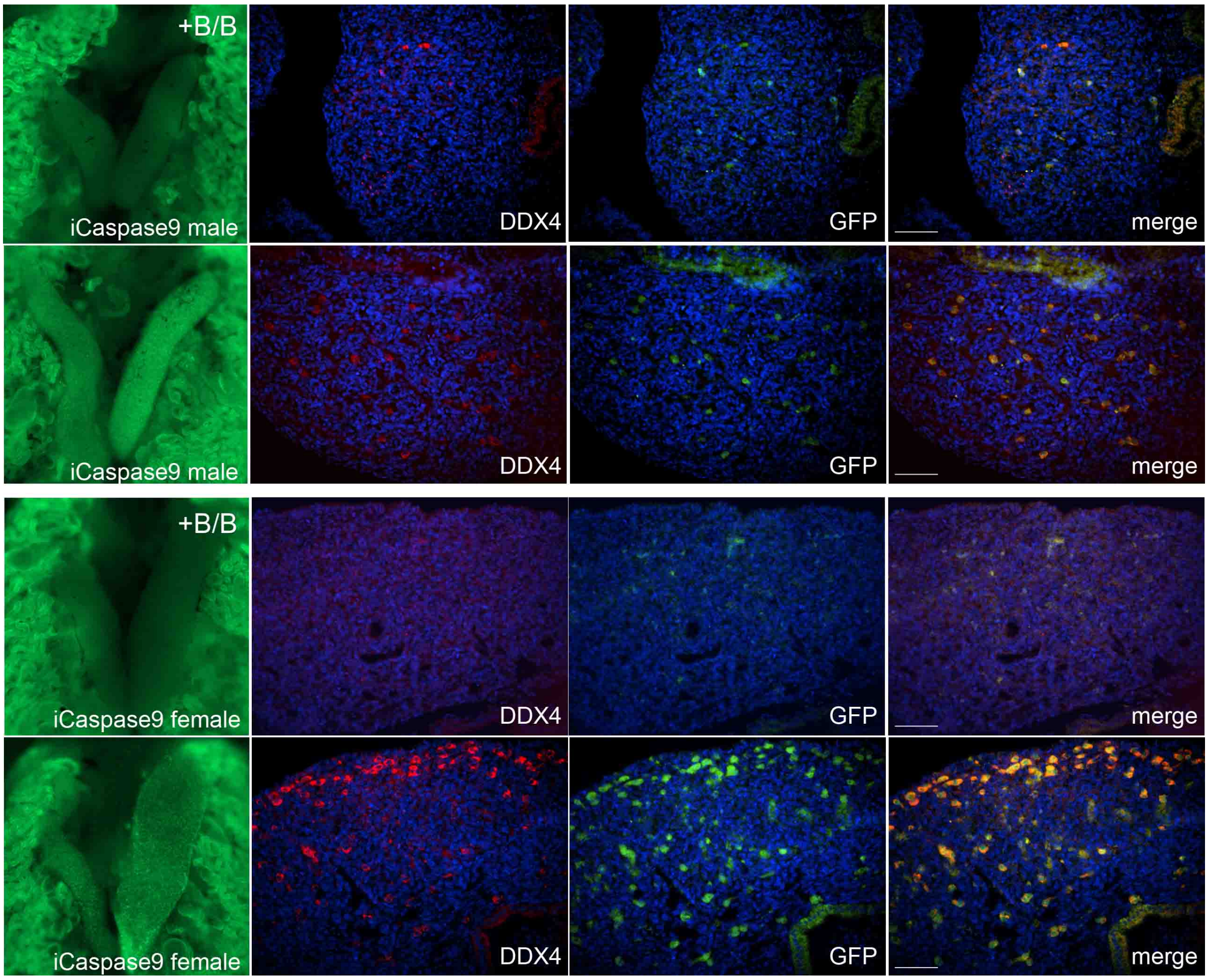

**Supplementary Fig. 6 GFP expression is germ cell specific in day 10 iCaspas9 G2 embryos and B/B treatment ablates endogenous PGCs**

B/B (1ul of 0.5mM B/B) was injected into the dorsal aorta of stage 16 chicken embryos (day 2.5). Embryos were incubated and examined at day 10 for GFP fluorescence (green) and DDX4 immunofluorescence (red). Size marker, 50 µm.

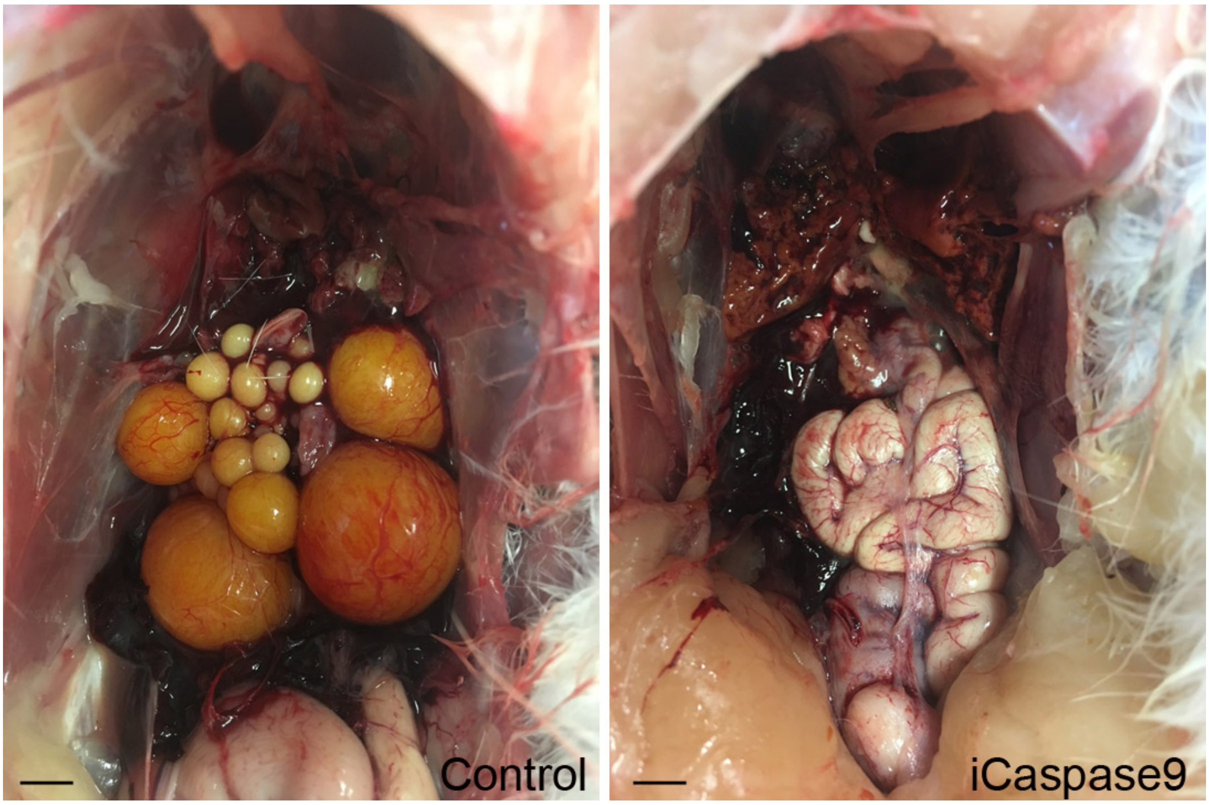

**Supplementary Fig. 7 Oocyte formation in B/B treated iCaspase9 female embryos**

Control and iCaspase9 embryos were microinjected with B/B compound (1 ul of 0.5mM) into the dorsal aorta at stage 16 (day 2.5). Embryos were incubated to hatch, hatched and raised to 31 weeks. The control hen contains a normal follicular hierarchy. The iCaspase9 hen contains no follicles. Scale bar, 1cm.

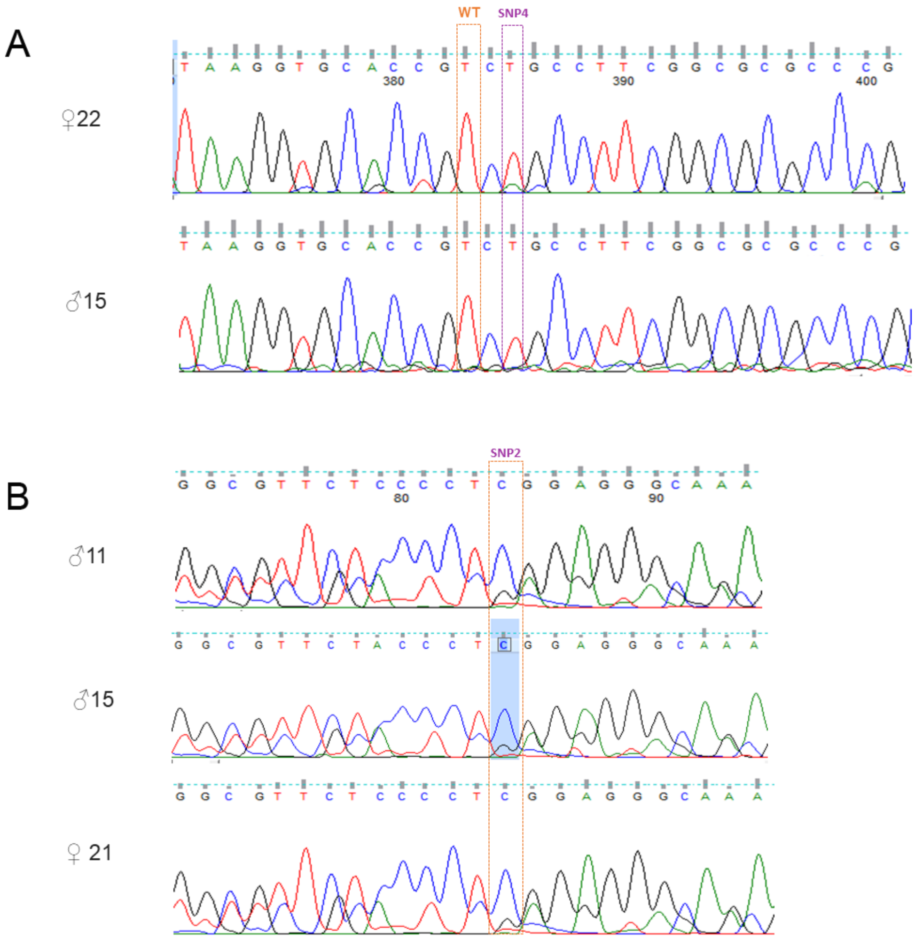

**Supplementary Figure 8 The B/B Dilution Barred feather allele is present in WL line 6 chicken.**

Screening of individual Line 6 chickens for mutations in the Z chromosome *cCDKN2A* gene **a,** homozygous for missense SNP4 R10C. **b,** homozygous for non-coding SNP2.

a

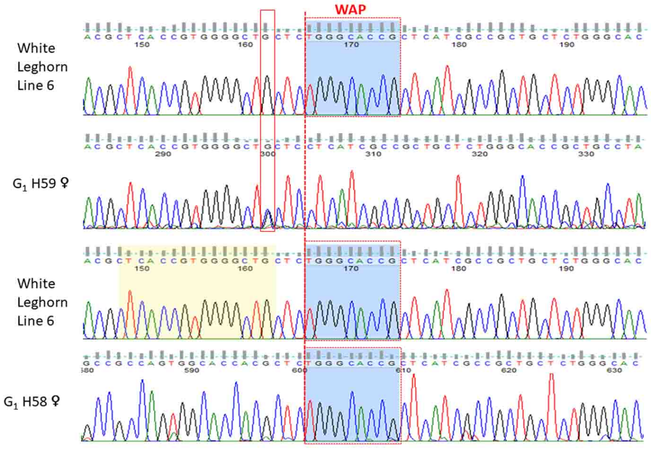

b

| Genotype | number | phenotype |
|---|---|---|
| *cPMEL17* $^{-WAP/-WAP}$ | 9 | 6 barred |
| | | 3 speckled |
| *cPMEL17* $^{-WAP/Del+WAP}$ | 8 | 7 barred |
| | | 1 speckled |
| *cPMEL17* $^{Del+WAP/Del+WAP}$ | 3 | 2 barred |
| | | 1 speckled |

c

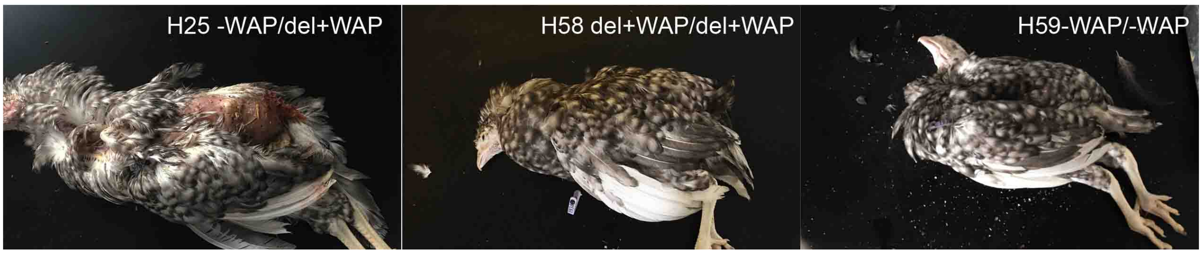

**Supplementary Fig. 9 Sequence and feather phenotypes of DOW edited $G_1$ birds**
**a,** Sequence of G1 offspring demonstrating the transmission of the edited pmel17 alleles. Bird H59 is homozygous for wildtype allele lacking the WAP insert, *cPMEL17* $^{-WAP/-WAP}$. The red box indicates the tracking allele introduced into female WL PGCs. Bird H58 is homozygous for two 15 bp deleted alleles.
**b,** Number of female offspring for all feather phenotypes. Female birds contained barred or specked feathers. The deleted allele was underrepresented in the hatched offspring however one of the surrogate male carried *cPMEL17* $^{-WAP/-WAP}$ edited PGCs and would not generate *cPMEL17* $^{\mathrm{del+WAP}}$ offspring.
**c,** Barred feathers in $G_1$ offspring. Similar feather colouring was present in all three genotypes.

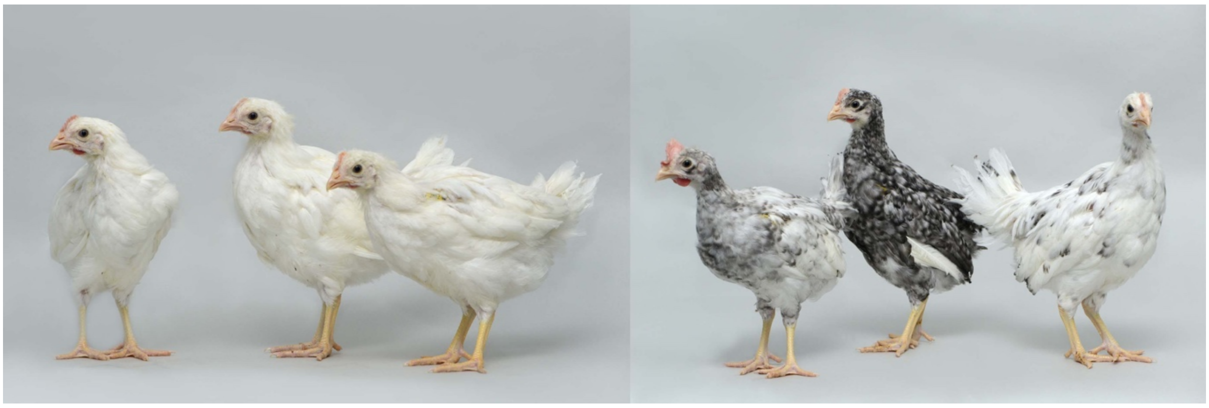

**Supplementary Fig. 10 G2 offspring from G1 DOW edited birds**

**a,** $G_2$ offspring from $G_1$H24♂ mated to Line 6 control females

**b,** $G_2$ offspring from $G_1$H24♂ mated to $G_1$H9♀ and $G_1$H12♀

### Supplementary Materials and Methods

#### iCaspase 9 vector design and construction

A 1.5 kb left targeting arm was PCR amplified using genomic DNA from Hy-line PGCs up to the *DAZL* stop codon. The middle fragment containing a 2a-GFP fragment was PCR amplified from an existing expression plasmid (ddx4-GFP). The right 1.5 kb targeting arm was PCR amplified from genomic DNA downstream from (and including) the *DAZL* stop codon. Gibson cloning was used to construct an initial Dazl left arm-2a\2a-GFP\Dazl right arm targeting vectors. Equimolar quantities of DNA fragments were incubated with the Gibson HiFi DNA Assembly Master Mix enzyme (NEB) for 2 hours at 50 °C and transformed into bacteria for plasmid isolation and sequencing.
The human and chicken 2A-iCasp9 sequences were commercially synthesised (Invitrogen) with BamH1 restriction sites using plasmid pMSCV-F-del Casp9.IRES.GFP (Addgene) as a template and subcloned into the DAZL-GFP construct (see supplementary materials and methods).

#### Guide design and CRISPR/Cas9 vector construction

CRISPR gDNAs were designed using CRISPOR gDNA design web tool (http://crispor.tefor.net) and CHOPCHOP gDNA design web tool. gDNA oligonucleotides were synthesised by Integrated DNA Technologies (IDT) and cloned into HF-PX459 V2.0 as previously described in [43]. The HF-PX459 V2.0 vector was used to express SpCas9-HF1 and sgRNA. Construction of the HF-PX459 V2.0 vector is previously described in [10]. All ssODN donors used were Ultramer DNA Oligonucleotides synthesised by IDT. ssODN and gRNA sequences are listed in Supplementary table 1. Microhomology-mediated end joining at the Pmel17 guide double strand break site were predicted using inDelphi and RGEN Tools microhomology predictors [29,30].

#### Detection of genome editing events

To detect editing events genomic PCR was performed with corresponding specific primer sets (see Supplementary Table 2). For *cKRT75* screening of PGC clones, cells were lysed in QuickExtractDNA extraction solution (Lucigen) following manufactures instructions. 2 µl of this lysate or 100 µg of gDNA were used as PCR template for *cKRT75* PCR amplification with High-Fidelity Phusion DNA polymerase (NEB) according to manufacturer's instructions. For PMEL screening, an initial PCR was performed using 200 ng of genomic DNA as PCR template for amplification with FastStart Taq DNA Polymerase (Sigma-Aldrich) according to manufacturer's instructions. Subsequently, to identify an indel in *PMEL17* indel another PCR was performed, 100 ng of genomic DNA was used a template for amplification with PrimerSTAR GXL DNA polymerase (Takara Bio) according to manufacturer's instructions. All PCR products were directly sequenced by Sanger sequencing to detect edited genotypes. Digestion of the PCR products with Pfi1 detected the *cPMEL17*$^{del}$ allele.

#### PCA plot

Genomic DNA was prepared from blood samples from $G_1$ chicks using cell lysis solution (Quiagen) containing RNAse A Solution (Sigma-Aldrich). Protein Precipitation Solution (Qiagen) was added and DNA was precipitated and resuspended. DNA from $G_1$ offspring and control (Hy-line Brown, iCaspase9, LSX, and Line 6) chicks were genotyped using a custom Cobb 60K Infinium Illumina array.

**Western blot analysis of expression of HA and GFP in CIA gonadal PGC**

Gonads from iCaspase9 or aviCaspase9 embryos at 14 days of incubation were examined for expression of GFP protein under fluorescent microscopy. The $GFP^+$ gonads were dissected and GFP-negative gonads were used as control. The gonads were lysed into RIPA buffer containing proteinase inhibitor. Protein was resolved on 5-14% SDS-PAGE gel (Bio-rad) and trans-blotted onto PVDF membrane. Anti-HA tag (Cell Signalling Tech.) and anti-GFP (Millipore) antibodies were used to probe the target proteins, anti-alpha tubulin antibody was used as loading control. To enrich for germ cells, gonads were also dissociated in parallel with collagenase/dispase enzyme (Sigma-Aldrich) and the cells were cultured in FAOT medium for 24 h. The cells in suspension were transferred to a new well on 24-well plate and further cultured for and additional 4 days. Suspension cells were harvested into RIPA buffer and used for western blots.

**Oligonucleotides:**

| Used for editing | Description | Sequence | length |
|---|---|---|---|
| Male Line 6 PGC line | DOW male repair template | GCATCCCCAGCCGCCAGTGGCACCACGCTCACCGTGGGGCTGCTCCTCATCGCCGCTGCTCTGGGCACCGCTGCCTACACCTACCGGTGAGCGGG | 95 bp |
| Female Line 6 PGC line | DOW female repair template | GCATCCCCAGCCGCCAGTGGCACCACGCTCACCGTGGGGCTCCTCCTCATCGCCGCTGCTCTGGGCACCGCTGCCTACACCTACCGGTGAGCGGG | 95 bp |
| Female LSX PGC line | Frizzle template | TCCCCAGCTCCCTCATCCTTTCTCCATAGGAGAAGGTTTTCCTGGAAGAGTCAGCCCGGCTGCGGTTGGCAATGTCCTCATACTGCGCCTTCACCTCGGC | 100 bp |
| Line 6 PGCs | PMEL17 guide | GCTCACCGTGGGGCTGCTCT | |
| Hy-line PGCs | DAZL guide | GGCTTACTAAACTGAACTGT | |
| LSX PGCs | cKRT75 guide | GGCTTCAGCCCGGCTGCGGT | |

PMEL targeting in female line 6 PGCs the HDR donor template was redesigned to include a 1 bp synonymous change. The change was introduced within 2 bp of the Cas9 cut site and also introduces a novel restriction site, Eco24I (BanII), see underlined in table above.

**PCR Primers:**

| Target | Primer name | Sequence | Product length |
|---|---|---|---|
| cPMEL17 | Pmel_exon10fwd | TGGCTGTGGCCAGCACCCACG | 382 bp |
| | MM-532 | GCAAACGCAGGGTAGCAC | |
| cKRT75 | cKRT75_F1 | AAGGCAGATGCATTGACCGA | 657 bp |
| | cKRT75_R1 | CATCAACTGCCCAGGGACTC | |
| iCaspase9 screening | mm786 | TGCCTGGTTGCTTTAATTTCCTC | 1017 bp |
| | mm787 | TGGAACAGGTAAAACAGAACACA | |
| aviCaspase9 screening | mm785 | GTCGACGGTGTCTCTGTGAA | 1813 bp |
| | mm787 | TGGAACAGGTAAAACAGAACACA | |
| Recessive white locus | Diag05-nor-up | CAAAACCATAAATAGCACTGGAAATAG | 481 bp |
| | Diag05-dw | TTGAGATACTGGAGGTCTTTAGAAATG | |
| | Diag05-cc-up | CCTCTGGCTCTATTTGACTACACAGT | 345 bp |
| | Diag05-dw | TTGAGATACTGGAGGTCTTTAGAAATG | |
| cGAPDH | c-gapdhF | CAGATCAGTTTCTATCAGC | |
| | c-gapdhR | TGTGACTTCAATGGTGACA | 700 bp |
| GFP | GFP-F | ACGTAAACGGCCACAAGTTC | 187 bp |
| | GFP-R | AAGTCGTGCTGCTTCATGTG | |

**DNA synthesised for human iCaspase9 transgene**
(2a-iCaspase9-Bam H1 site mutated)
GGATCCGGAGCTACTAACTTCAGCCTGCTGAAGCAGGCTGGAGACGTGGAGGAG
AACCCTGGACCTATGCTCGAGGGAGTGCAGGTGGAGACTATCTCCCCAGGAGAC
GGGCGCACCTTCCCCAAGCGCGGCCAGACCTGCGTGGTGCACTACACCGGGATG
CTTGAAGATGGAAAGAAAGTTGATTCCTCCCGGGACAGAAACAAGCCCTTTAAG
TTTATGCTAGGCAAGCAGGAGGTGATCCGAGGCTGGGAAGAAGGGGTTGCCCAG
ATGAGTGTGGGTCAGAGAGCCAAACTGACTATATCTCCAGATTATGCCTATGGTG
CCACTGGGCACCCAGGCATCATCCCACCACATGCCACTCTCGTCTTCGATGTGGA
GCTTCTAAAACTGGAATCTGGCGGTGGTTCCGGAGTCGACGGATTTGGTGATGTC
GGTGCTCTTGAGAGTTTGAGGGGAAATGCAGATTTGGCTTACATCCTGAGCATGG
AGCCCTGTGGCCACTGCCTCATTATCAACAATGTGAACTTCTGCCGTGAGTCCGG
GCTCCGCACCCGCACTGGCTCCAACATCGACTGTGAGAAGTTGCGGCGTCGCTTC
TCCTCGCTGCATTTCATGGTGGAGGTGAAGGGCGACCTGACTGCCAAGAAAATG
GTGCTGGCTTTGCTGGAGCTGGCGCGGCAGGACCACGGTGCTCTGGACTGCTGCG
TGGTGGTCATTCTCTCTCACGGCTGTCAGGCCAGCCACCTGCAGTTCCCAGGGGC
TGTCTACGGCACAGATGGATGCCCTGTGTCGGTCGAGAAGATTGTGAACATCTTC
AATGGGACCAGCTGCCCCAGCCTGGGAGGGAAGCCCAAGCTCTTTTTCATCCAG
GCCTGTGGTGGGGAGCAGAAAGACCATGGGTTTGAGGTGGCCTCCACTTCCCCT
GAAGACGAGTCCCCTGGCAGTAACCCCGAGCCAGATGCCACCCCGTTCCAGGAA
GGTTTGAGGACCTTCGACCAGCTGGACGCCATATCTAGTTTGCCCACACCCAGTG
ACATCTTTGTGTCCTACTCTACTTTCCCAGGTTTTGTTTCCTGGAGGGACCCCAAG

AGTGGCTCCTGGTACGTTGAGACCCTGGACGACATCTTTGAGCAGTGGGCTCACT
CTGAAGACCTGCAGTCCCTCCTGCTTAGGGTCGCTAATGCTGTTTCGGTGAAAGG
GATTTATAAACAGATGCCTGGTTGCTTTAATTTCCTCCGGAAAAAACTTTTCTTTA
AAACATCAGTCGACTATCCGTACGACGTACCAGACTACGCACTCGACCTCGACG
GATCC

**DNA synthesised for chicken iCaspase9 transgene**
(2a-iCaspase9)
AGGGATCCGGAGCTACTAACTTCAGCCTGCTGAAGCAGGCTGGAGACGTGGAGG
AGAACCCTGGACCTATGCTCGAGGGAGTGCAGGTGGAGACTATCTCCCCAGGAG
ACGGGCGCACCTTCCCCAAGCGCGGCCAGACCTGCGTGGTGCACTACACCGGGA
TGCTTGAAGATGGAAAGAAAGTTGATTCCTCCCGGGACAGAAACAAGCCCTTTA
AGTTTATGCTTGGCAAGCAGGAGGTGATCCGAGGCTGGGAAGAAGGGGTTGCCC
AGATGAGTGTGGGTCAGAGAGCCAAACTGACTATATCTCCAGATTATGCCTATG
GTGCCACTGGGCACCCAGGCATCATCCCACCACATGCCACTCTCGTCTTCGATGT
GGAGCTTCTAAAACTGGAATCTGGCGGTGGTTCCGGAGTCGACGGTGTCTCTGTG
AATTGCAGACCAGCTAGGATGCATGCTAGTGCATGCCAGGTGTACCAGCTGCGA
GCAGACCCTTGTGGGCACTGCCTGATCTTCAACAATGTCAGCTTCAGCAGAGACT
CTGATCTGTCGACTCGAGCTGGCTCTGACATAGACTGTGAGAAGCTGGAGAAGC
GTTTCAGGTCCCTGTGCTTCCACGTCCGGACCCTGCGGAACCTCAAAGCTCAGGA
AATTGATGTGGAGCTGCGGAAGCTGGCGCGGCTCGACCACAGTGCCCTGGACTG
CTGCCTCGTGGTCATCCTCTCCCATGGTTGCCAGACAAGCCATATTCAGTTTCCCG
GAGGGATTTATGGAACAGATGGCAAAATCATTCCAATCGAAAGGATTGTGAACT
ATTTCAATGGGTCCCAGTGCCCGAGTTTGAGAGGAAAACCCAAACTCTTCTTCAT
CCAGGCCTGTGGAGGAGAACAAAAGGACCAAGGATTTGAGGTGGATTGTGAATC
ACCCCAAGATGAAACTTGCCGACGTTCCATAGAGTCGGATGCGATTCCTTTCCAG
GCTCCATCAGGGAATGAGGACGAGCCAGACGCCGTCGCCAGTTTGCCCACTCCT
GGTGACATCTTGGTGTCCTATTCAACTTTTCCAGGTTTTGTGTCCTGGAGGGACA
AGGTGAGTGGCTCGTGGTACGTGGAAACCTTGGACAGCGTACTGGAACATTACG
CCCGTTCTGAAGACCTGCTTACCATGCTACTTCGGGTGTCAGACATCGTATCCAC
CAAGGGGAGGTACAAGCAGATCCCGGGCTGTTTCAACTTCCTTCGTAAAAAATTC
TTCTTCCTGTGCAAGGTCGACTATCCGTACGACGTACCAGACTACGCACTCGACG
GATCCAA
